# Biomolecular AFM viewer

**DOI:** 10.1101/827402

**Authors:** Romain Amyot, Holger Flechsig

## Abstract

We developed a stand-alone software which allows to transform biomolecular structures and movies of their conformational dynamics into a representation which corresponds to the outcome of biomolecular atomic force microscopy (AFM) experiments, such as high-speed AFM. The software implements a high degree of usability. An integrated highly versatile molecular viewer allows the visualization of structures and their corresponding simulated AFM representations in arbitrary orientations. The obtained results can be conveniently exported as still images and movies. We provide a demonstration of our biomolecular AFM viewer by applying it to several proteins from the Protein Data Bank, and to a molecular movie of conformational transitions between two protein structures obtained from a modelling server.

## Introduction

*Seeing is believing* paraphrases the holy grail of molecular biophysics - the imaging of molecular processes which establish Life at the nanoscale. At the single protein level this task is particularly challenging, due to the tininess of structures and the rapid timescales of functional conformational motions. Large progress has been made by the development of the high-speed atomic force microscopy (hs-AFM) technique, which overcomes many limitations inherent to traditional single-molecule approaches such as fluorescent resonance spectroscopy. Hs-AFM allows to observe conformational dynamics in proteins under physiological conditions at high spatio-temporal resolution. ^1,2^

The power of hs-AFM experiments was first demonstrated by monitoring the operation of the bacteriorhodopsin protein^3^ and the important motor proteins myosin V^4^ and F1-ATPase.^5^ To date, hs-AFM is a leading method to observe dynamical processes in proteins and its success has been evidenced in a plethora of studies. ^6^

Like any other experimental approach hs-AFM has also limitations. The temporal resolution is strictly limited by the video rate with which *protein movies* can be recorded (typically less than 10 frames per second). The spatial resolution allows to resolve well motions of domains but is typically not high enough to visualize changes of individual structural elements which may be of functional importance. A third drawback is the control of the molecular orientation upon depositing the proteins on the surface. Furthermore, in a single experiment observation proceeds from a fixed viewpoint, generally limiting the information on the coupling of motions between opposite regions within a protein.

By improving technological aspects of hs-AFM, those drawbacks can be overcome only up to a certain extent. In fact, a promising strategy to further advance our understanding may consist in a complementary approach, combining experiments with available structural data and in particular computational modelling of biomolecular systems.

The three-dimensional structure of most biological macromolecules is known to date, available from different experimental methods (X-ray diffraction, nuclear magnetic resonance, electron microscopy). This data is deposited in a single global archive, the Protein Data Bank (PDB),^7,8^ and provides the spatial coordinates of atomic positions in a well-defined format (the PDB file). Although this data captures a protein in a static frozen conformation, its great availability offers valuable reference information for the understanding of dynamic hs-AFM experiments.

On the other side, structural data provides the basis for computational modelling of protein dynamics. Multi-scale modelling constitutes a standard approach of computational biophysics to explore functional motions in proteins at different levels of complexity. Often simulations are performed using automatized software packages and require costs manageable by usual PC workstations. In particular, coarse-grained models allow to record *molecular movies* of proteins which capture the functional conformational motions during many operation cycles. ^9,10^ A comparison between *computational molecular movies* obtained from simulations with *molecular movies* recorded in hs-AFM experiments becomes thus important. This requires the development of suitable computational tools. We report here a standalone software which allows to transform the three-dimensional molecular structure of a protein into a graphical representation which corresponds to the output of AFM experiments, referred by us to as a simulated AFM image. The software serves two major tasks

- transforming an experimentally known static protein structure, provided by a PDB-file, into its simulated AFM image
- processing molecular movies of protein dynamics (in PDB format) to produce simulated AFM movies.

The software is designed to have no dependencies on any commercial products and implements a high degree of usability. An integrated highly versatile molecular viewer allows the visualization of molecular structures and their corresponding simulated AFM analogues, for both static structures and molecular movies. In a *what-you-see-is what-you-get* like fashion, results can be conveniently exported as image files and also movies.

We will first explain the molecular viewer as the framework underlying our software. We then describe the procedure developed to produce simulated AFM images from PDB structural data. In the application part of this paper we give examples which demonstrate the comparison of both PDB structural and movie data to hs-AFM experiments of proteins.

## Molecular viewer of protein structures and movies

The software can process static structures and molecular movies as long as they are provided in the standard PDB file format. To load a PDB file its corresponding ID has to be provided. For structures available from the PDB archive the information will be processed online. In a PDB file of a molecular movie the ATOM record containing the spatial coordinates have to be embedded into the MODEL and ENDMDL environment.

The software integrates a molecular viewer to visualize the three-dimensional structure of a protein as encoded in the corresponding PDB file. All atoms provided in the PDB file are displayed in the viewer window. Two representations are available. In the *VdW representation* all atoms are displayed as spheres with the radii each given by the specific Van der Waals (VdW) radius of the atom type. In the *surface representation* a smoothing algorithm with variable resolution is applied to visualize the shape of the protein.

The orientation of the loaded protein structure can be arbitrarily changed through rotations and zooming within the viewer window.

## Simulated AFM images

In the hs-AFM experiment of biomolecules the output represents the height of the sample surface. This information is then processed to produce graphical images of the sample shape in which the height is color-coded.

We describe how to transform the protein structure into a simulated AFM image. The underlying procedure translates the set of spatial coordinates of atoms into a corresponding height map, taking into account the tip shape and protein representation. The method we apply relies on the tip-structure collision condition which has been previously employed. ^5,11,12^ It is illustrated in Fig. 1. The tip is assumed to have a cone-like shape with a probe sphere at its narrow end. Its geometry is characterized by the cone half-angle *α* and the probe sphere radius *R* (see Fig. 1A). To determine the height map of the protein in a given orientation, the corresponding molecular structure is scanned by the tip over a two-dimensional grid with the spacing parameter *a* (see Fig. 1B). For each cell within this grid the height is determined from the collision condition of the tip shape with the VdW spheres of all atoms. For a detailed mathematical formulation of this procedure we refer to the Supplementary Information.

**Figure 1:**
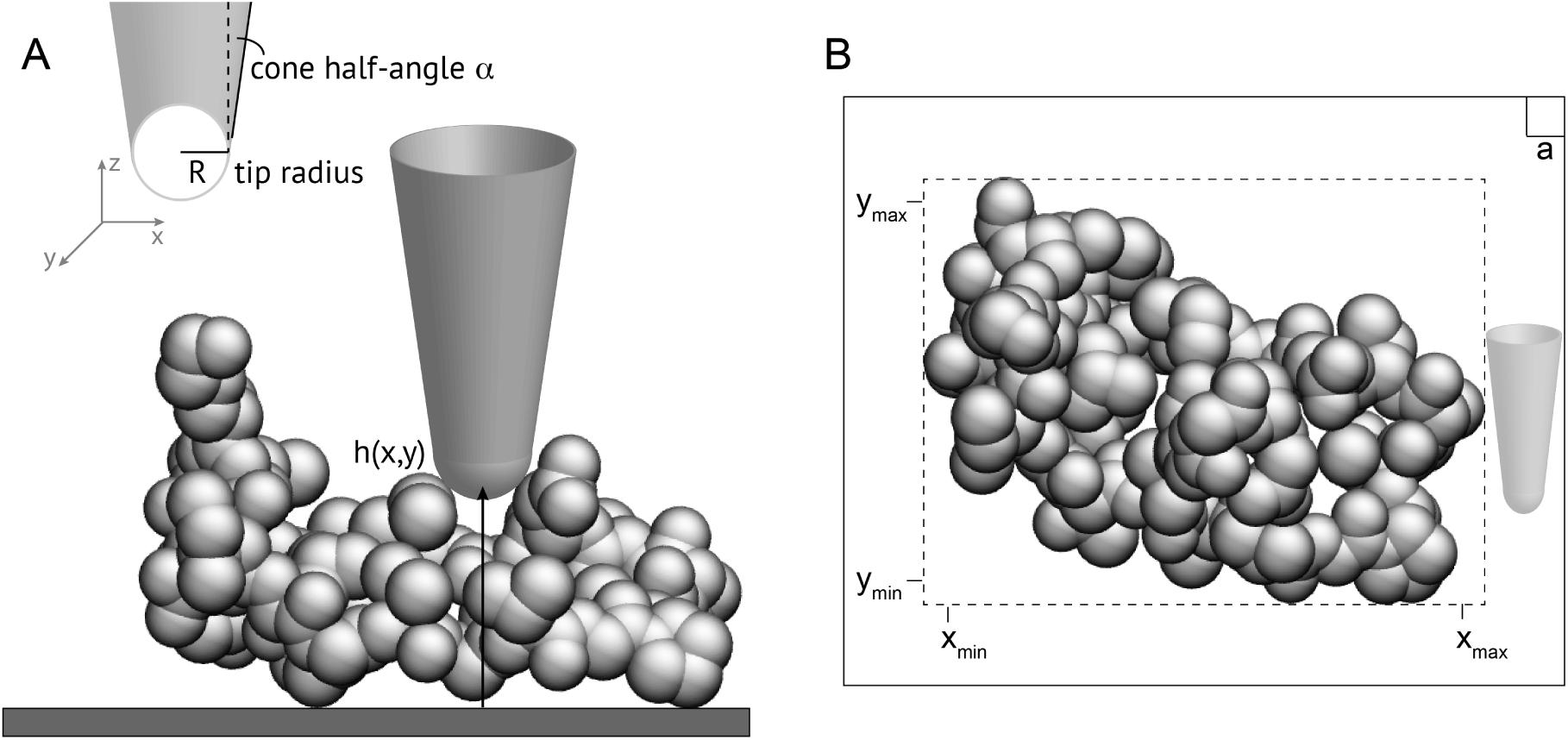
The cone-structure collision method to generate simulated AFM images. A, the tip shape and the collision condition of the tip with the protein VdW structure is illustrated. B, the size of the scanning area is shown with the spacing parameter *a* of the imposed grid.

## Demonstration of the AFM viewer

To provide a demonstration of the AFM viewer we have chosen the structure of the G-actin protein (PDB ID 3HBT) and that of the membrane transporter MsbA (PDB ID 3B60, chain A and B). The structure of the G-actin protein is shown in Figure 2 in the all-atom VdW representation and in the surface representation, as they are displayed in the viewer window of our software. The corresponding simulated AFM images are also provided.

**Figure 2:**
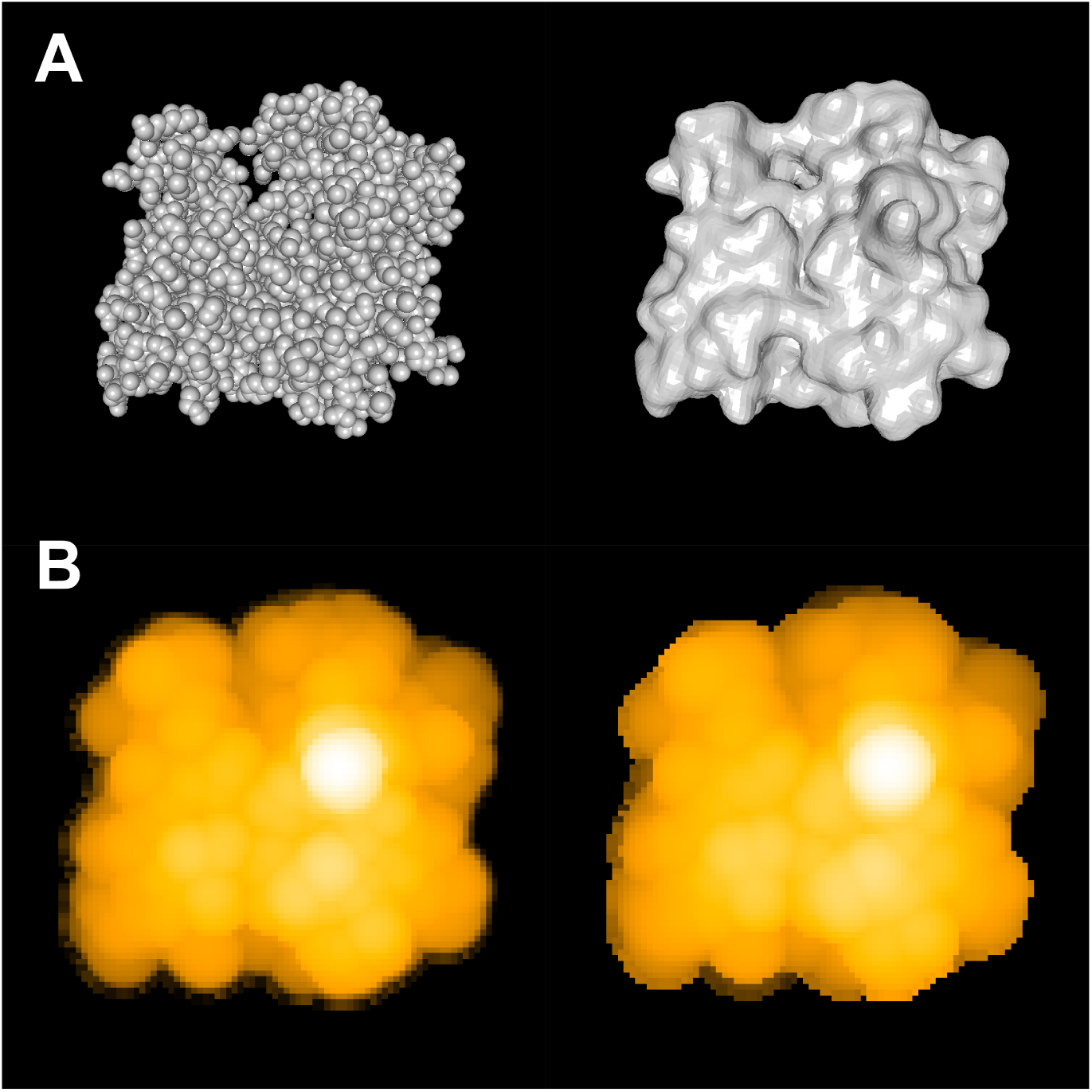
Demonstration of the AFM viewer. A) The top row shows the structure of G-Actin as displayed in the viewer window. The all-atom VdW representation and the surface representation is shown. B) In the bottom row the corresponding simulated AFM images as displayed in the viewer window are shown. The hypothetically chosen parameters were *a* = 1Å, *R* = 5Å and *α* = 5°.

In Figure 3 the MsbA transporter is shown in the all-atom VdW representation as displayed in the viewer window. The structure of the transporter is shown in three different orientations with their corresponding simulated AFM images.

**Figure 3:**
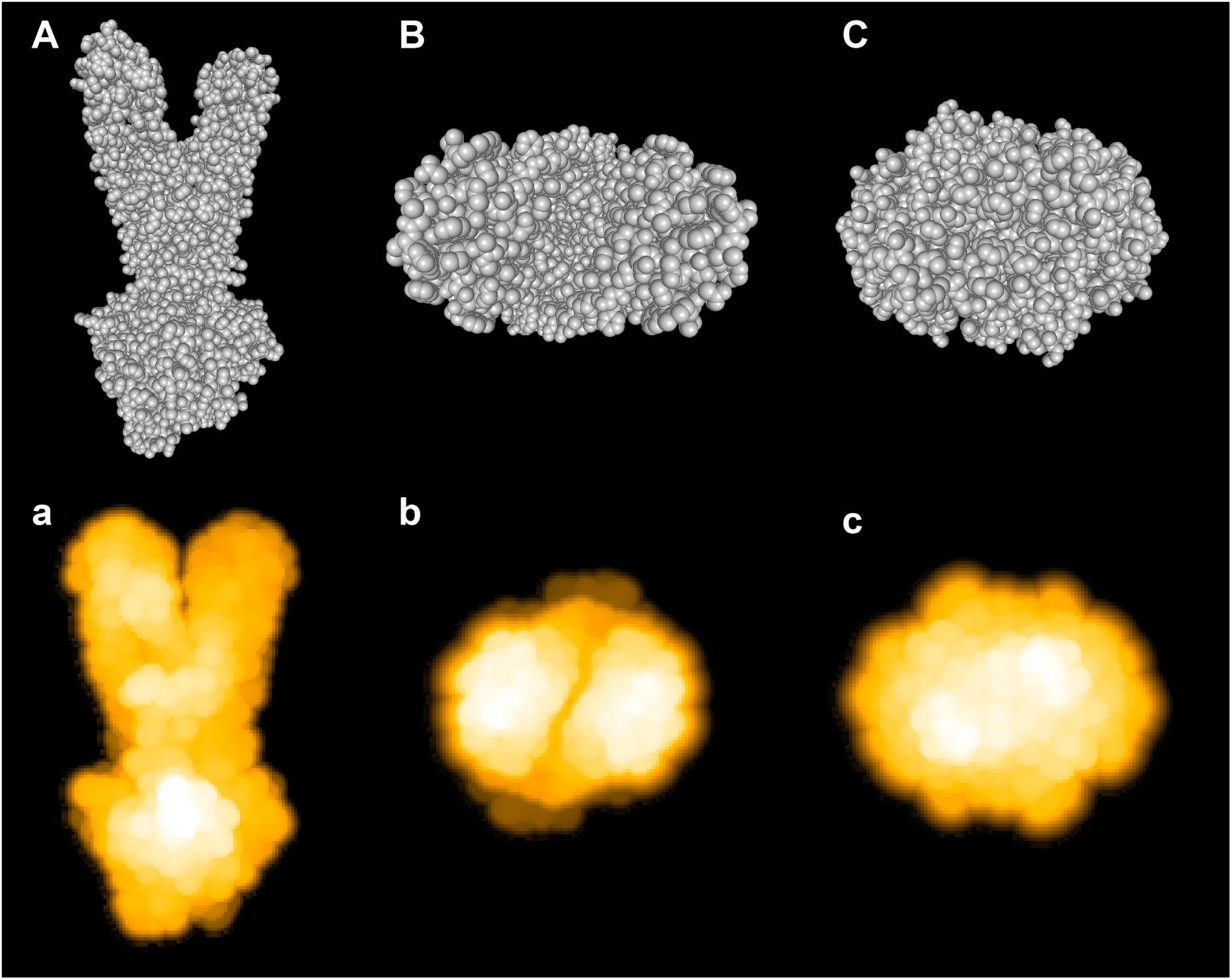
Demonstration of the AFM viewer. Top row: the static structure of the MsbA transporter (PDB ID 3B60, chain A and B) as displayed in the integrated molecular viewer using the Van der Waals representation. Different orientations are shown: A, the front view. B, the top view showing the trans-membrane domains. C, the bottom view showing the nucleotide-binding domains. Bottom row: the corresponding simulated AFM images are shown. The hypothetically chosen parameters were *a* = 1Å, *R* = 3Å and *α* = 5°. We note that the molecular structures (A,B,C) are shown in the perspective view.

For the purpose of demonstration we have chosen hypothetical parameters to construct the simulated AFM images. When a comparison with hs-AFM experimental images becomes available, the parameter values can be adjusted accordingly.

### Effect of tip shape on AFM image resolution

To demonstrate the effect of the tip geometry on simulated AFM images we have considered the structure of the actomyosin complex known from the PDB structure in a fixed orientation. In Figure 4 we show simulated AFM images which have been constructed for different values of the cone-half angle *α* and different values of the probe sphere radius *R*. For the purpose of demonstration we have used hypothetical parameters. We understand though that they are beyond the reach of current experimental realization. We observe two effects, both expected as a result of the applied tip-structure collision. First, an increase in the probe radius *R* leads to a larger coarsening in the resolution of the molecular structure. Second, an increase in the cone half-angle lead to a larger blurring, visible at the boundaries of the protein structure. When the step-size is increased the images get more pixelated (see Fig. 4, bottom row). This effect is typically present in hs-AFM images of biomolecules caused by the limitations in the spatial resolution.

**Figure 4:**
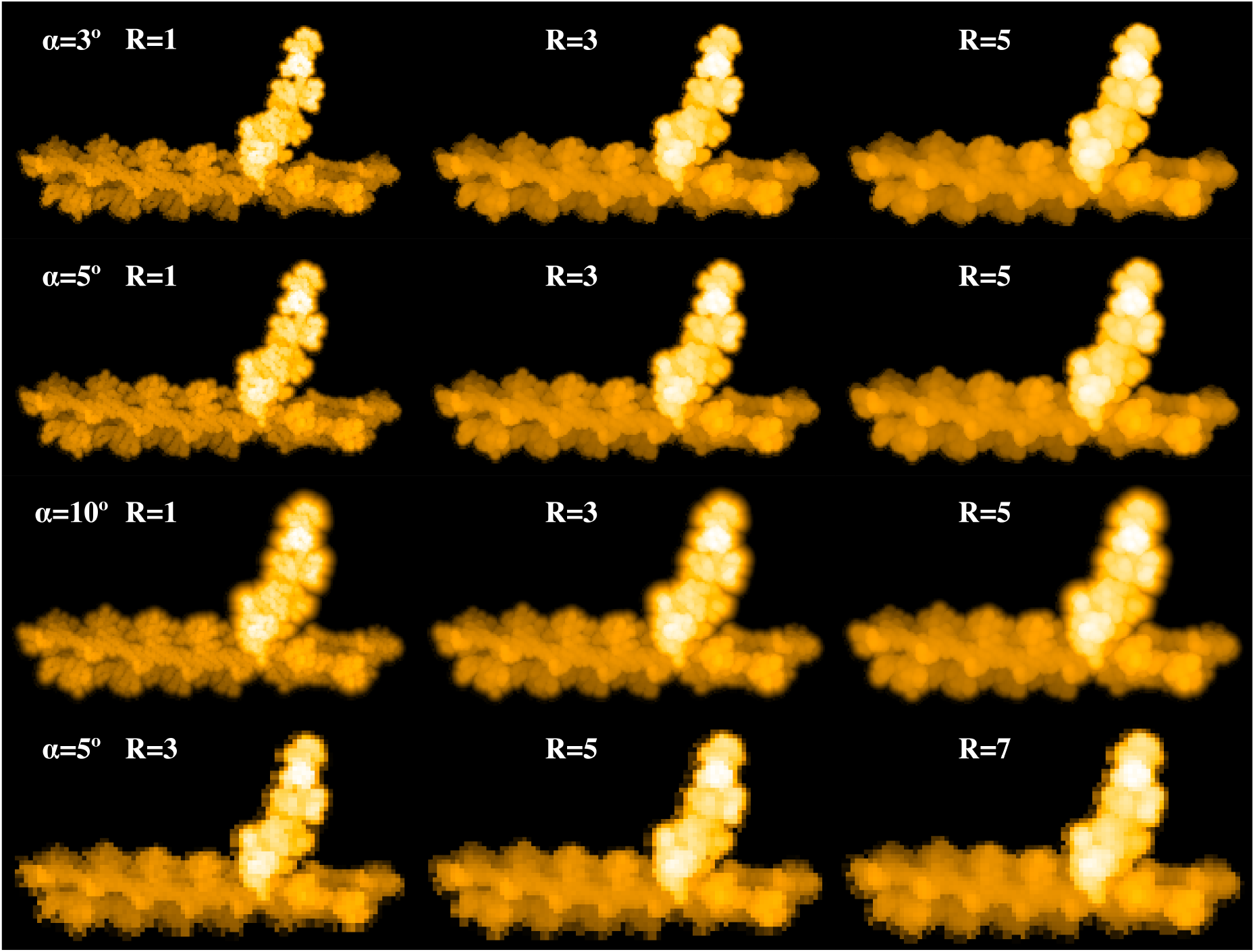
Effect of cone shape geometry in simulated AFM images. The structure of the actomyosin complex (PDB ID 1M8Q, myosin chains P,Q,R) is shown as simulated AFM images using different values for the cone half-angle *α* and the probe sphere radius *R* (in Å). Images in the first three rows were obtained with a hypothetical spacing parameter of *a* = 2Å for scanning. For the bottom row *a* = 5Å was used. When generating simulated AFM images, only the alpha-carbon atoms of the structure were considered here.

### Molecular movie of conformational transition

The program can load a molecular movie of arbitrary origin, provided it is stored in the PDB file format. It is required that the ATOM record of consecutive frames is embedded into the MODEL and ENDMDL environment, which corresponds to the output format of standard modelling servers. After loading, the movie of conformational changes will be displayed in the viewer window which has the same versatile functionality as described before. After setting the AFM parameters, the corresponding simulated AFM movie will be generated. For the purpose of demonstration we have considered a molecular movie of the open to close transition in the GroEL monomer (structures of PDB ID 1SX4 were used). The feasible conformational transition pathway was computed by the iMODS server based on the normal mode analysis.^13^ In Figure 5 we show snapshots of the movie taken at different time moments in the VdW representation, and we show the corresponding snapshots of the generated simulated AFM movie.

**Figure 5:**
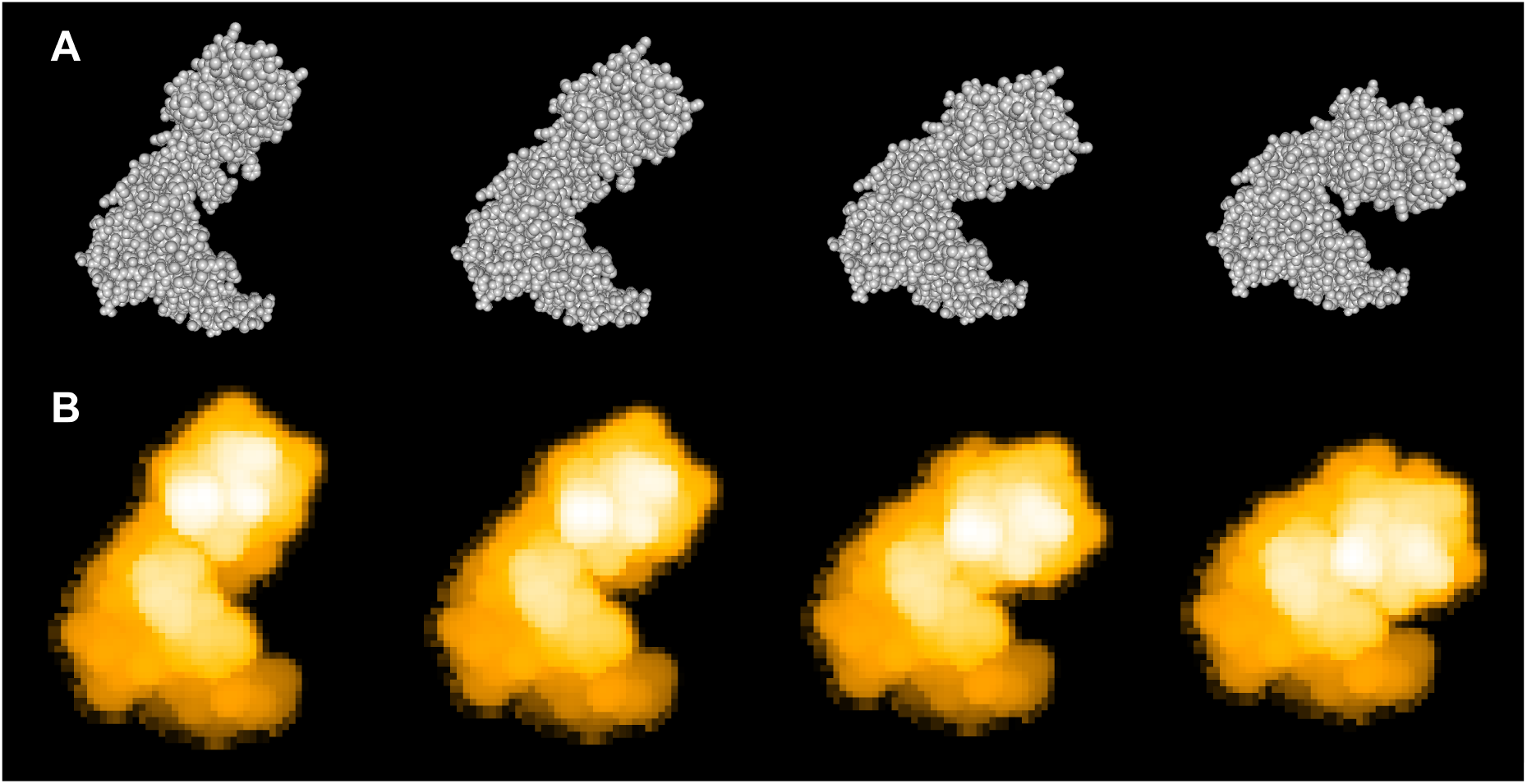
Molecular movie of GroEL. A) The top row shows snapshots of the open to close transition in the GroEL monomer. B) Corresponding snapshots of the simulated AFM movie. We have used hypothetical parameters *a* = 2Å, *α* = 5° and *R* = 5Å for scanning.

### Exporting images and movies

The program allows to take snapshots of the graphics displayed in the viewer window by a single click. They will be stored as image files in different possible file formats. We have also implemented a function which generates rotations of the content shown in the viewer window. On demand the graphics in the viewer window can be exported as a movie which records all performed action. The file will be stored in different possible movie formats.

## Conclusions

We have developed a stand-alone software which allows to transform biomolecular structures and movies of their conformational dynamics into a representation which corresponds to the outcome of biomolecular atomic force microscopy (AFM) experiments, such as high-speed AFM. An integrated highly versatile molecular viewer allows the visualization of structures and their corresponding simulated AFM representations in arbitrary orientations. The obtained results can be conveniently exported as still images and movies. Here we have provided a first demonstration of this biomolecular AFM viewer by applying it to several proteins from the Protein Data Bank. We have also demonstrated its application to a molecular movie of conformational transitions between two protein structures obtained from a modelling server.

## Author contributions

H.F. has designed the study, prepared the figures and wrote the manuscript. R.A. has developed the software.

## Acknowledgement

We thank Noriyuki Kodera and Kien Xuan Ngo from the Nano Life Science Institute at Kanazawa university for explaining the hs-AFM experiments to us.

## Supporting Information Available

We provide details on how to generate simulated AFM images as a supplementary text.

## Software availability

We will make the developed software available for use on Linux OS and the Windows operating system.

