## Supplementary text for "Biomolecular AFM viewer"

### 1 Shape of the tip

The tip is composed of a cone characterized by a cone half-angle $\alpha$ and a sphere of radius $r$. The sphere is placed into the cone in such a way that the centre $S$ of the sphere is aligned to the vertex $V$ of the cone and that the sphere intersects with the cone along a circle (see Figure 1). The cone region below the sphere (see shaded area in Figure 1) does not contribute to the tip shape. To calculate the collision between the tip and a spherical atom, we calculate both the collision between the spherical part of the tip and the sphere of the atom, and the collision between the conic part of the tip and the spherical atom. In the latter case, however, we discard collisions with the shaded region and the atom sphere.

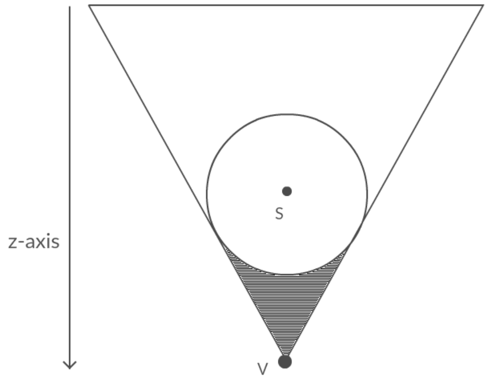


Figure 1: Schematics of the probe shape.

To determine the height of the atomic structure needed to generate simulated AFM images, we have to compute the hard collision of the tip shape with the atom spheres.

### 2 Collision between a cone and a sphere

We consider a cone with a vertex $V = (x_V, y_V, z_V)$ and a sphere of centre $S = (x_S, y_S, z_S)$ and radius $r$ (see Figure 2). A point $(x, y, z)$ on the surface of the cone is separated by a distance $a$ from the central axis. The projection of this point to this axis is separated by a distance $c$ from the vertex. We have $c = z_V - z$ and $a = (z_V - z)\tan(\alpha)$.
A point $(x, y, z)$ on the surface of the sphere is separated by a distance $b$ from the central axis and the projection of this point to this axis is separated by a distance $d$ from the centre. We have $d = (z_S - z)$ and $b = \sqrt{r^2 - (z_S - z)^2}$.
We also consider the two dimensional distance $f$ between the vertex $V$ and the centre $S$, $f = \sqrt{(x_V - x_S)^2 + (y_V - y_S)^2}$.
For the collision point $I = (x, y, z)$, the collision condition is fulfilled (see Figure

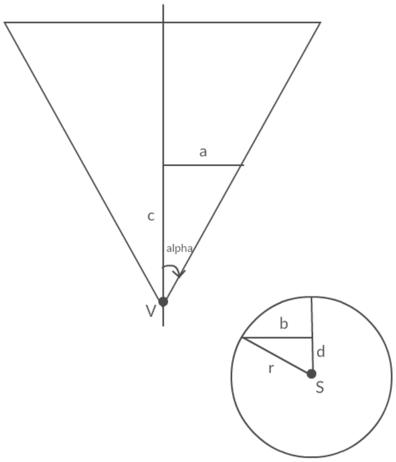


Figure 2: Schematics of cone and sphere with the used distances indicated.

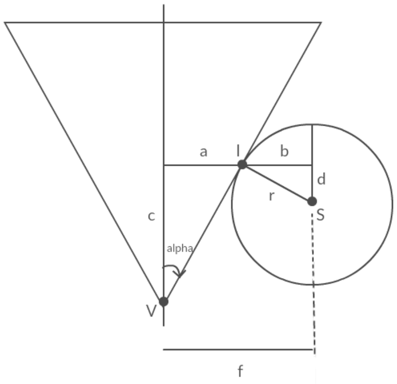


Figure 3: Schematics of a collision between the cone and the sphere.

3). It reads

$$a + b = f. \tag{1}$$

We have $(z_V - z)\tan(\alpha) + \sqrt{r^2 - (z_S - z)^2} = f$,

which leads to a second-order polynomial equation in $z$:

$$(1 + \tan^2(\alpha))z^2 + (-2z_V \tan^2(\alpha) + 2f\tan(\alpha) - 2z_S)z \\ + (f^2 - 2fz_V \tan(\alpha) + z_V^2 \tan^2(\alpha) - r^2 + z_S^2) = 0 \tag{2}$$

Depending on $z_V$ this equation can have either no solution, i.e. no intersection (Figure 2), one unique solution(Figure 3), or 2 solutions (Figure 4).
The unique solution case corresponds to the collision where the cone is tangent to the sphere.

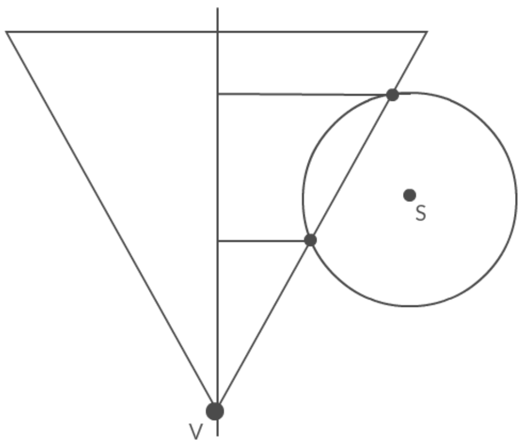


Figure 4: Schematic representation of the sphere penetrating the cone, see text for explanation.

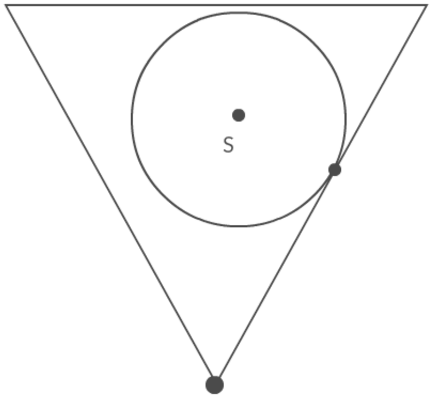


Figure 5: Schematics of cone and sphere collision, which is discarded. See text for explanation.

This corresponds to the vanishing discriminant $\Delta$ of the polynomial in equation (2), which will determine $z_V$. This gives a second order polynomial equation in $z_V$

$$(-\tan^2(\alpha))z_V^2 + (2z_S\tan^2(\alpha) + 2f\tan(\alpha))z_V + (-2fz_S\tan(\alpha) - f^2 + r^2 + r^2\tan^2(\alpha) - z_S^2\tan^2(\alpha)) = 0. \quad (3)$$

The discriminant of the corresponding polynomial is $\Delta_2 = 4r^2\tan^2(\alpha)(\tan^2(\alpha)+1) > 0$.

Then, equation (3) has always two solutions. It means there are two positions of the vertex for which the equation (2) has a unique solution. They are shown in Figure 3 and Figure 5. The smaller solution $z_V$ is taken.

The solutions are $z_V = \dfrac{z_S \tan(\alpha) + f \pm r\sqrt{\tan^2(\alpha) + 1}}{\tan(\alpha)}$
and $z = \dfrac{z_V \tan^2(\alpha) - f \tan(\alpha) + z_S}{1 + \tan^2(\alpha)}$.

### 3 Collision between two spheres

A sphere $S = (x_S, y_S, z_S)$ of radius $r$ collides with a sphere $P = (x_P, y_P, z)$ of radius $r_P$ (the probe sphere) if the sum of their radii is equal to the distance between their centres.
The collision condition is

$$f^2 + (z_S - z)^2 = (r + r_P)^2,$$

where $f = \sqrt{(x_P - x_S)^2 + (y_P - y_S)^2}$.

This given a second-order polynomial equation in $z$,

$$z^2 - 2z_s z + (f^2 - (r + r_P)^2 + z_S^2) = 0.$$

The discriminant is $\Delta = -4(f^2 - (r + r_P)^2)$.

The solutions are $z = z_S \pm \sqrt{(r + r_P)^2 - f^2}$. The smaller solution is taken.
